# Dynamic Graphs Analysis of EEG

**DOI:** 10.1101/2025.07.30.667711

**Authors:** Mohamed Radwan, Pedro G. Lind, Anis Yazidi

**Affiliations:** Oslo Metropolitan University, Department of Computer Science, 0167 Oslo, Norway; Kristiania University of Applied Sciences, School of Economics, Innovation and Technology, 0107 Oslo, Norway; Simula Research Laboratory, Numerical Analysis and Scientific Computing, 0164 Oslo, Norway; University of Oslo, Department of Informatics, 0373 Oslo, Norway

**Keywords:** Graphs, EEG, Dynamic Connectivity, GNNs

## Abstract

In this study, we investigate the use of temporal dynamics in brain connectivity for the classification of electroencephalography (EEG) signals using dynamic Graph Neural Networks (GNNs). Our methods are applied to several large-scale EEG datasets focused on abnormality and epilepsy detection. The implemented models demonstrate competitive performance on unseen test subjects across all three datasets, outperforming previous graph-based baselines in terms of accuracy and F1 score. We explore multiple architectures designed to capture temporal variations in graph-structured data, demonstrating their effectiveness in modeling dynamic brain activity. In addition to classification, we employ graph-theoretical metrics to analyze temporal changes in brain networks, such as network efficiency and node degree, across time windows of EEG recordings. The goal is to characterize differences between pathological and healthy groups at both the node and network levels. We particularly examine epilepsy and healthy subject groups to highlight differences in local network efficiency and node degrees, with statistical significance confirmed via F-tests.

## 1 Introduction: Scope and background

Electroencephalography (EEG) has significant potential as a diagnostic tool for neurodegenerative diseases. It offers the advantage of capturing brain dynamics and activity with high temporal resolution. Various techniques have been employed to analyze the EEG features for the purpose of predicting neurodegenerative diseases. However, understanding the brain dynamics responsible for the neurodegenerative diseases remains an open research question. Furthermore, analysis of EEG data is a challenging task given the complexity of features. For example, studying one feature alone can be misleading. The brain dynamics is mostly a combination of several EEG features that can be helpful to understand the diseases. We aim from this study to provide a simple and comprehensive approach that shows how to integrate dynamic features in the analysis of EEG.

Neural networks are widely utilized for EEG analysis. In particular, Graph Neural Networks (GNNs) have gained attention for their ability to incorporate the geometrical and topological relationships between EEG electrodes into the modeling process. In this article, we focus on GNNs that operate on dynamically evolving graphs. These dynamic graphs represent time-varying relationships among brain regions and we aim to employ those dynamics in predictive modeling tasks.

EEG signals are recorded from multiple electrodes placed on the scalp, and the signals can be modeled as a graph where the nodes correspond to EEG channels (electrodes), and the edges represent functional connectivity between them. Various functional connectivity measures can be used to define these edges, including Coherence, Mutual Information, and Phase Locking Value (PLV), among others.

A dynamic graph is one in which the connectivity (edges) and, in some cases, the nodes change over time. Such representations reflect how functional brain connectivity varies during a cognitive task or in response to a stimulus. The use of dynamic graphs in EEG analysis is vital because the brain is essentially non-static. Functional interactions between brain regions can change rapidly, and modeling the brain with static graphs may oversimplify this complexity. Dynamic graph models allow us to capture of temporal variations in connectivity, thereby providing richer and more informative features for neural networks.

Several trials using GNNs were developed on time evolving graphs. All the methods are based on combining graph to capture the geometrical features of the graph with sequence based models to capture the temporal dynamics. These methods are developed in other domains such as traffic forecasting. These models includes EvolveGCN [17], Diffusion Convolutional Recurrent Neural Network (DCRNN) [12], Temporal Graph Convolutional Network (TGCN) [28], Attention temporal graph convolutional network (A3T-GCN) [1], Graph Convolutional Recurrent Networks [21], Graph convolution embedded LSTM (GC-LSTM)[2], and ROLAND framework [26].

We notice that the use of dynamic Graphs Neural Networks in EEG is not well explored in research. One method is the use of temporal GNNs are used for Seizure Detection [6] is used utilizes the features the temporal and spatial dimensions in the EEG data. The model uses several spatiotemporal convolutional layers. The study uses Short term Fourier Transform (STFT) that are manually extracted spectrogram features for each time segment to be used as node features. This study ignores the temporal correlation between channels. This means that the spatial coupling between electrodes are fixed through the whole EEG graph. Another study used DCRNN from [12] on EEG data [23] for seizure prediction utilizing self supervision in predicting the next EEG window taking current window which proved to provide superior performances. The study [22] uses Spatial-temporal graph convolutional network for Alzheimer classification. The authors uses different connectivity measures Pearson Correlation, Coherence, Phase Locking value and Phase lag index. This network consists of two blocks which composes of temporal 1D-Conv and Spatial Graph Convolution. However, this study ignores the temporal connectivity changes between EEG nodes.

The study [5] developed InstaGAT based on Graph Attention Networks and combined it with LSTM for EEG classification. The model is used on TUAB dataset [16]. This model serves as a good candidate baseline for comparison. The study [29] developed Stacked GAT-LSTM for classification of EEG. The model is mainly LSTM layers that processes the sequence of GAT embedded vectors. The model is used on TUAB dataset [16]. Similarly, this model serves as a good baseline for comparison.

Our contributions in this study are:

- Using temporal graph neural networks for the classification of EEG. The model utilizes the dynamic topology of graphs using the time evolving adjacency matrices.
- Provide a parameter efficient baseline that utilize the advanced temporal dynamics in the graph based on the temporal geometrical operators such as DCRNN, TGCN, AT3-GCN, GConvGRU and GCLSTM. These operators are explained in details in section 2.6.
- Report the significance of connectivity evolving dynamics between the groups of subjects from EEG data. We apply this method for epilepsy disease.

The details about this study starts with section 2 which gives the details about the used data, preprocessing, experimental settings and the architecture of the model used. Section 3 report the results achieved by the used models and further analysis of the dynamic graphs between the groups of data subjects. Finally, in the discussion section 4, we resort to previous research to validate the findings in this study and conclude the limitations.

## 2 Data and Methods

### 2.1 Dataset

We use three datasets in this study for different prediction tasks. The first dataset is the TUH Abnormal dataset (TUAB) [16], which consists of 2993 EEG records. The dataset has 2717 train subjects and 276 test subjects. The test set has 127 abnormal and 150 normal subjects. The meaning of Abnormal is that the EEG contains clear evidence of pathology as interpreted by a neurologist. Several Graph models such as [5] and [29] are applied on this data which we use as baselines for our evaluation of this study.

The second dataset is the TUH Epilepsy Corpus (TUEP) which is a subset of the Temple University Hospital EEG Corpus (TUH EEG Corpus) [16]. The dataset has 200 Unique subjects. It is hard to evaluate the model on this dataset given that the test data is not predefined and chosen randomly. This would make unfair comparisons. The study by [15] analysed several static functional connectivity and used convolutional neural networks for classifications.

The third dataset is the NMT Scalp EEG dataset [10], which consists of 2417 EEG records. The EEG montage used for recordings is the standard 10-20 system. The dataset has 1972 normal and 445 abnormal subjects. The abnormal here means that there is a problem in an area of brain activity which are diagnosed to various neurological conditions. The test set size is 185 subjects with 95 normal and 90 abnormal subjects.

### 2.2 Experiments

In our experiments, we measure the metrics: Accuracy and F1-score for evaluation. The source code for the experiments is available at: https://github.com/mhmdrdwn/dgraph. TUAB and NMT datasets have predefined test holdout set. The TUEP dataset is split randomly into 90% train and 10% test subjects. Cross validation and hyperparameter tuning are done on the train data. The test data for all the used datasets are holdout unseen subjects to be used for the final evaluation of the used model. All the reported evaluation metrics is subject based classification. The models are used for binary classification to predict whether the entire EEG is normal or abnormal.

### 2.3 Preprocessing

The data is filtered using bandpass filter (1:45) Hz for all the datasets. All the datasets are downsampled using sampling frequency of 100 Hz. Each signal in broken down to windows. Each window has the size of two seconds or 200 time points. The number of windows are 200, 60 and 100 for the TUAB, TUEP and NMT datasets, respectively. Those windows are used to calculate the functional connectivity between EEG electrodes. The methods for computing the functional connectivity is explained in details in section 2.4.

### 2.4 Functional Connectivity

EEG raw signals windows are used to calculate Coherence and Phase Locking Values (PLV) functional connectivity. Each connectivity is computed for each window of EEG signals, leaving us to a dynamic graph over time. The size of the windows is two seconds. The choice of window size is not straightforward. There are several studies to analyze the effect of the window size on the computations of features. The study by [14] used fusion of different windows calculated to improve the performance on fMRI dataset. In our study, we only used two seconds of windows due to the expensive runtime of computing the connectivity. As explained earlier, we extract connectivity measures for each time window for each subject. This leaves us with connectivity tensor of shape *n, n, t* for each EEG where *t* is the number of windows and *n* is the number of nodes or EEG channels. The following connectivity measures are used in this study.

**Coherence** is a frequency-domain measure to assess the functional connectivity between two signals, particularly in neurophysiological studies involving electroencephalography (EEG). It quantifies the degree of linear correlation between two signals as a function of frequency. It is used to assess the synchronization between brain regions.

Given two time series signals *x*(*t*) and *y*(*t*), the coherence function *C*_*xy*_(*f*) at frequency *f* is defined as

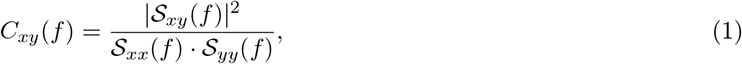

where:

- 𝒮_*xy*_(*f*) is the cross-spectral density between *x* and *y*,
- 𝒮_*xx*_(*f*) and *𝒮*_*yy*_(*f*) are the auto-spectral densities of *x* and *y*, respectively.

The coherence value ranges between 0 and 1, where 0 indicates no linear relationship at frequency *f*, and 1 denotes a perfect linear relationship.

#### Phase Locking value

PLV [11] is a phase-based measure of functional connectivity used to quantify the consistency of phase differences between two signals or strength of synchronization between different brain regions across multiple trials or over time. It is particularly useful in EEG and MEG studies for detecting phase synchronization between brain regions. PLV is commonly used to It is particularly valuable because it is robust to amplitude fluctuations and focuses solely on phase relationships. Given two signals *x*(*t*) and *y*(*t*), their instantaneous phases *ϕ*_*x*_(*t*) and *ϕ*_*y*_(*t*) can be extracted using the Hilbert transform or wavelet transform. The PLV is defined as

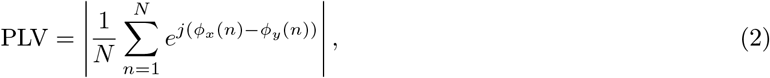

where:

- *N* is the number of time points or trials,
- *j* is the imaginary unit,
- *ϕ*_*x*_(*n*) and *ϕ*_*y*_(*n*) are the instantaneous phases of signals *x* and *y* at time point or trial *n*.

The PLV value lies in the range 0 and 1, where 1 indicates perfect phase locking and 0 indicates random phase differences.

### 2.5 Used Architecture

The EEG signals are split into windows of two seconds size with no overlap before the calculation of connectivity measures such as PLV and Coherence. This leaves a window of 100 time points given the sampling frequency 100 Hz. So, the size of those windows signals need to be reduced. This will be explained in figure 2. The reduced size embedding is used along with the connectivity matrices and fed to the graph temporal operators such as TGCN, AT3GCN, DCRNN, GConvGRU and GC-LSTM. The theory behind those operators are explained in section 2.6. The outputs of those operators are hidden states of the EEG nodes. Global Mean Pooling and Fully Connected layer are applied on those hidden states to get the output of the model where softmax is applied to predict the class.

**Fig. 1:**
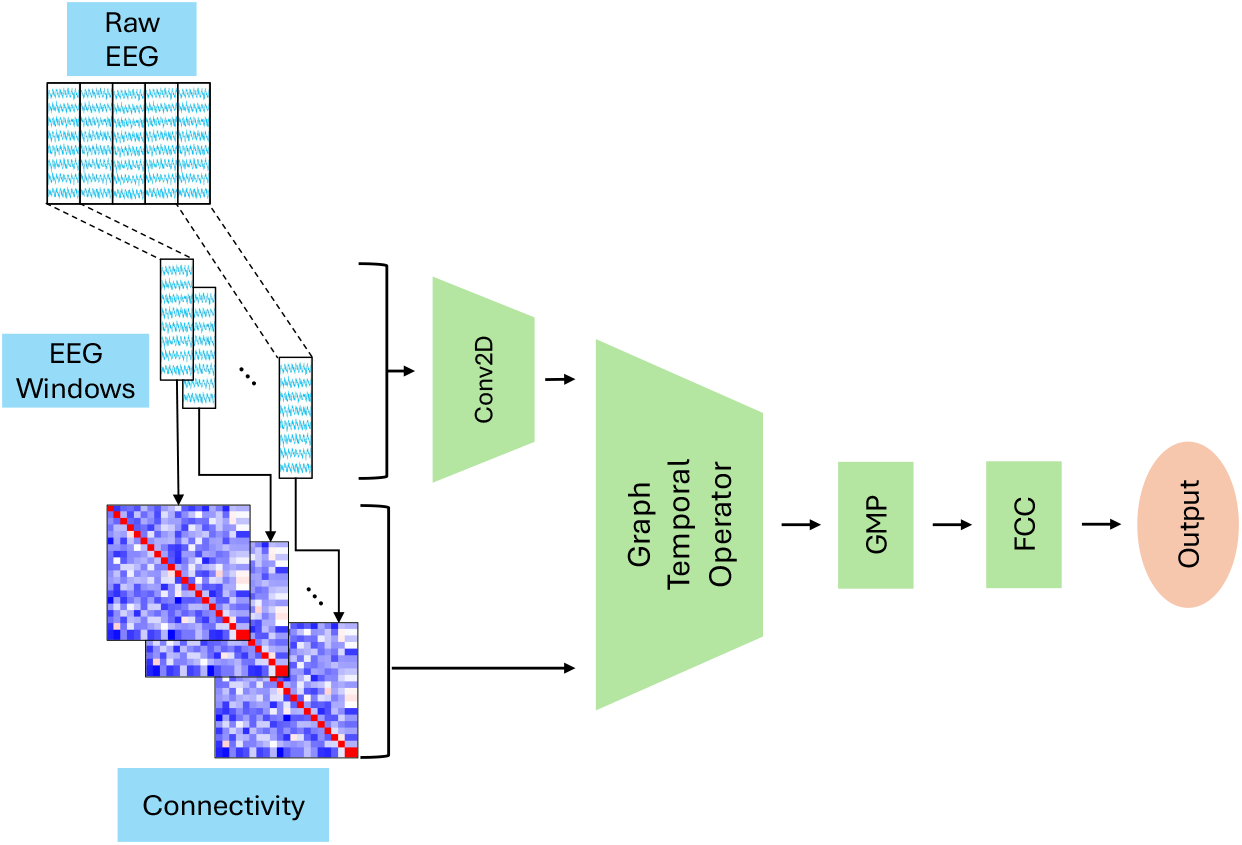
Used Predictive Architecture. The Raw EEG data is split into windows where the connectivity measures are calculated. The EEG windows are fed into Conv2D encoder 2 before fedding to the graph temporal operator. Finally, Global Mean Pooling (GMP) is applied on the output of the graph temporal operator before Fully Connected layer (FCC) to generate the output.

**Fig. 2:**
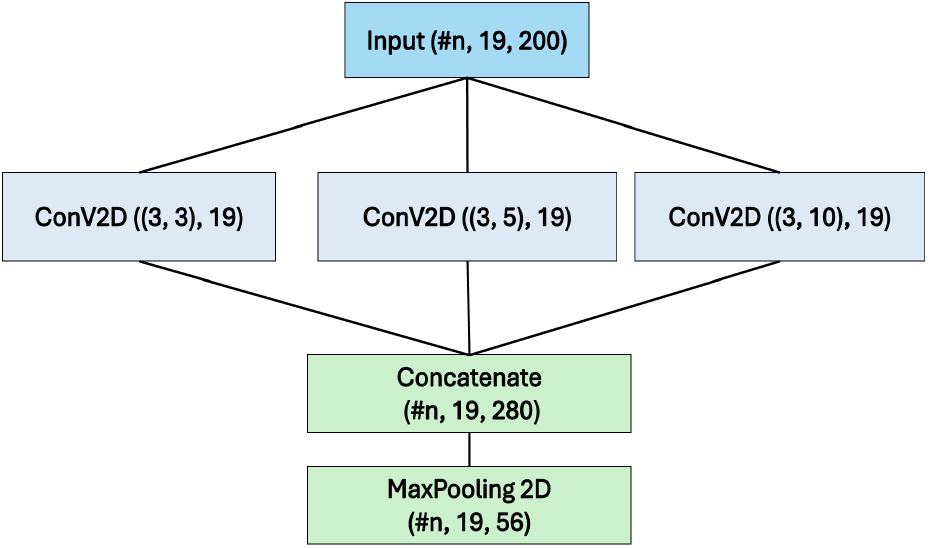
2D Convolutional Block to encode the time series windows before feeding to the graph operators. Three 2D convolutional filters are used wit different kernel sizes. Convolutional filters output is concatenated along the embedded time point features before feeding to Max Pooling layer to output the same number of channels as before.

As explained, the EEG data has extensive sequential details due to the long length of signals. the size of the windows is 100 time points. This makes training sequence models such as Recurrent Neural Networks difficult. To tackle this problem, we use Convolutions in addition to Maxpooling to reduce the size of the temporal sequence. Given the consecutive windows, a Convolutional Neural Networks (CNN) module is used to encode those windows before feeding to the GNNs models. The module architecture is shown in figure 2. The module is inspired by [18] which consists of several 1D Convolutional blocks with different filter sizes before concatenation. The objective from [18] is to reduce the size of the time series before feeding it to sequence model. Here, we use use a 2D Convolutional filters to reduce the size of the time points before feeding to the GNNs models.

### 2.6 Graph Temporal Operators

The Graph Temporal Operators are the core building block of the used architecture 1. It can be one of the following operators:

#### Temporal Graph Convolutional Network

TGCN [28] integrates Graph Convolutional Networks (GCNs) into the recurrent update equations of a Gated Recurrent Unit (GRU) to model spatiotemporal data. Each time point is convolved with GCN using the adjacency matrix and fed to GRU layers. The GRU captures the historical dynamics in the time points. At the end, the GRU output hidden state for after feeding each time point.

The Figure 3 show the network of TGCN. TGCN is mainly two steps model. It performs first Spatial Modeling where it applies Graph Convolutional Network (GCN) to extract spatial features from the graph. The second step is temporal modeling where it uses a Gated Recurrent Unit (GRU) to capture temporal dependencies over time. The spatial modeling is calculated using the following equation:

**Fig. 3:**
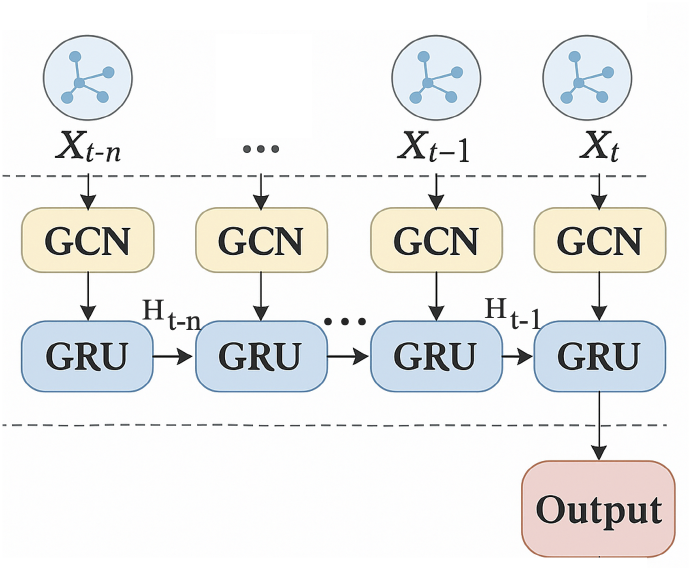
TGCN network (modified after [28]). The Learned Temporal features for each time stamp (X) and Adjacency matrices are fed into GCN and then fed to GRU along with the hidden state from previous time stamp (Shown as *H*_*t−n*_, *H*_*t−*1_).

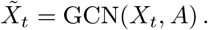

The GCN takes the features (Output of the window embedding) *X* along with adjacency matrix *A*. The GCN operation is defined as

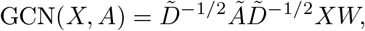

where *Ã* = *A* + *I* is the adjacency matrix with self-connections, 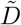 is the degree matrix of *Ã, W* is a learnable weight matrix. The temporal modeling is calculated using the following equation:

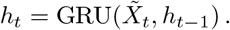

Here, GRU captures temporal patterns across sequences of spatial features.

#### Attention Temporal Graph Convolutional Network

A3T-GCN [1] extends TGCN [28] by incorporating attention mechanisms in the spatial and temporal dimensions. A3T-GCN attention is simply a scoring function (i.e. MLP) is designed to calculate the importance of each hidden state. In A3T-GCN, TGCN is used to output hidden states from the time windows. A scoring function is designed to calculate the score/weight of each hidden state. Third, an attention function is designed to calculate the context vector that can describe global temporal variation information. Let *H* = [*H*_*t−T*_, …, *H*_*t*_] be the sequence of the hidden states. The temporal attention score *α*_*t*_ is computed as

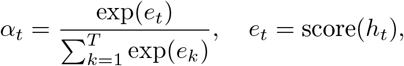

where *e*_*t*_ is the score of the hidden state at time *t*. This gives the context vector as

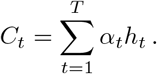

#### Graph convolution embedded LSTM

GC-LSTM [2] integrates Graph Convolutional Networks (GCNs) within the Long Short-Term Memory (LSTM) architecture to jointly model connectivity and temporal dependencies. This architecture is similar to previously mentioned TGCN [28] but replacing the GRU with Long Short Term Memory (LSTM) [9]. In TGCN, the memroy cell has only hidden states. Here, the memory cell of this network has cell states in addition to hidden states.

#### Graph Convolutional GRU

GConvGRU [21] replaces the linear operations in the GRU with Graph Convolution operations to model both spatial and temporal dependencies. Unlike TGCN [28] which is a two steps model, GConvGRU is one unified cell that simultaneously learns fusion connectivity and temporal dependencies interact. GConvGRU cell is an extension of GRU [4] which is used to update the hidden state with the most relevant information in the temporal sequence. This is achieved through gates that learn relevant sequential information. The GRU update gate, reset gate, candidate hidden state and final hidden state are defined as

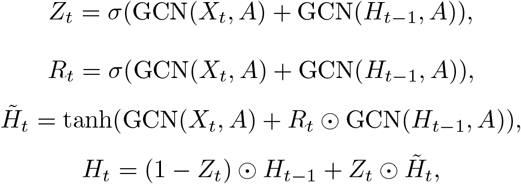

where *H*_*t−*1_ be the hidden state at time *t* −1, ⊙denote element-wise multiplication and *σ* be the sigmoid function. Here, we used the implementation provided by [19] that used Chebyshev convolutional operator [7] in all GRU convolutions.

#### Diffusion Convolutional Recurrent Neural Network

DCRNN [12] integrates a graph diffusion process with a recurrent neural network to model spatiotemporal data. The bidirectional diffusion convolution is defined is defined using a bidirectional random walk diffusion process:

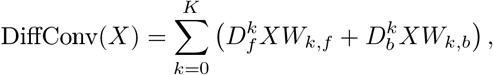

Here, *D*_*f*_, *D*_*b*_ be forward and backward diffusion matrices, *W*_*k,f*_, *W*_*k,b*_ be trainable weights for each diffusion step *k* and *K* is the number of diffusion steps. DCRNN is exactly similar to GConvGRU but with the graph diffusion DiffConv instead of GCN.

## 3 Results

In this section, we start by showing the basic analysis of the dynamic connectivity. Then we show the usage of dynamic connectivity in classification of EEG data using GNNs. First, we use the analysis of network efficiency, Node degree and their evolution over time of EEG signal from the epilepsy TUEP dataset. Then, we use the previously mentioned temporal graph operators based GNNs to classify the EEG on the three used datasets. Here we examine whether that the 60 snapshots contains the dynamic connectivity evolving features show patterns for the different subjects.

### 3.1 Network Efficiency

Efficiency is a graph-theoretical measure that quantifies how efficiently information is exchanged across the network. Here, we studied the Global efficiency and Local efficiency. We aim to study the dynamic changes in efficiency over time using the TUH Epilepsy Corpus (TUEP) [16]. As mentioned earlier, the TUEP dataset has two subject groups: healthy control and epilepsy subjects. Figures 4 and 5 shows the boxplots of the global and local efficiency of the subjects from the two groups.

**Fig. 4:**
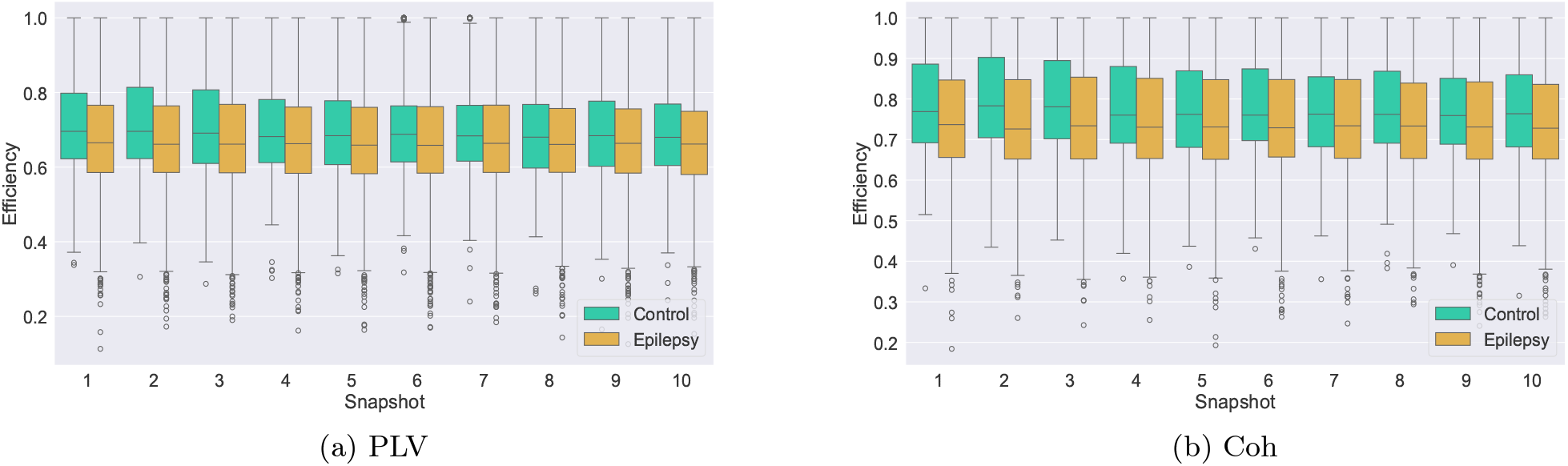
Dynamic Global Efficiency differences between the two groups of subjects: control vs epilepsy for a)Phase Locking Value (PLV) and b)Coherence (Coh).

**Fig. 5:**
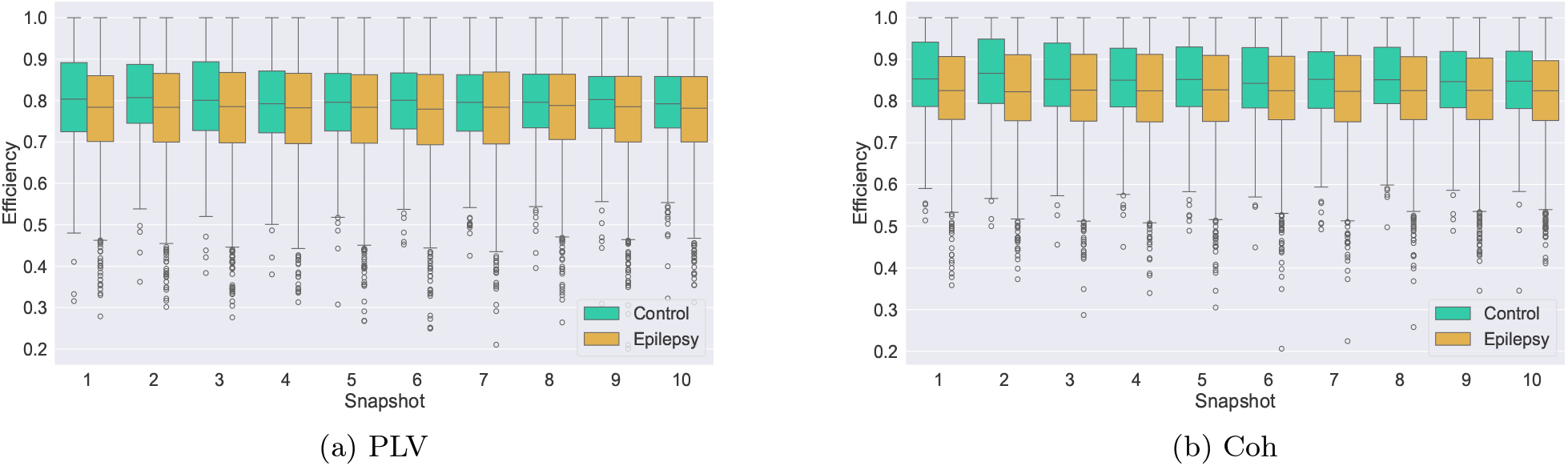
Dynamic Local Efficiency differences between the two groups of subjects: control vs epilepsy for a)Phase Locking Value (PLV) and b)Coherence (Coh).

Figure 4 shows the the group differences global efficiency between the epilepsy and control. Overall, the global efficiency is lower, in general, for epilepsy group. This difference is maintained through the time points or snapshots. Similarly, Figure 5 shows the local efficiency between the two groups. It is shown, in general, that the local efficiency is lower in epilepsy group. This difference is also maintained through the time points.

Figure 6 shows the dynamic fluctuations of global and local efficiency through time that differs for each subject. We studied the significance of the efficiency fluctuations for the two groups. The p-values for the efficiency fluctuations are: 0.00015 and 0.000054 for local Phase Locking Value and Coherence local efficiencies implying significance. However, we notice no significance in the global efficiency fluctuations between the two groups.

**Fig. 6:**
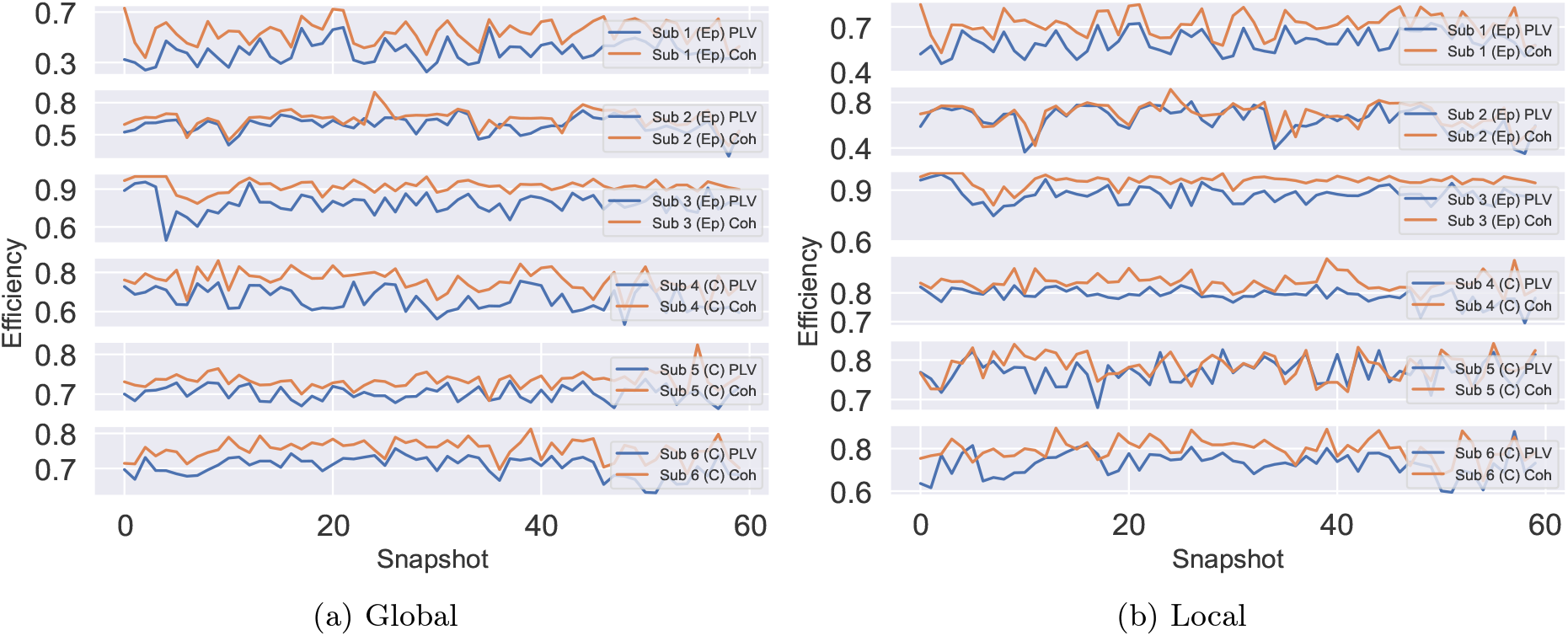
Dynamic Phase Locking Value and coherence efficiency for sample subjects from the two groups: control (C) vs epilepsy (Ep) for a)Global and b)Local Efficiency

### 3.2 Node Degree

Node degree represents the number of connections a node has in a network. Node degree here changes over time as the network evolves. This change can be observed as a result of addition or removal of edges over time. The degree of a node reflects its importance or centrality in the network. Nodes with high degrees are often referred to as hubs, playing crucial roles in the connectivity and dynamics of the network.

Figure 7 shows the dynamic evolution of node degrees for example different subjects from the two groups: epilepsy and control. The subfigure 7a shows the patterns for different subjects while the subfigure 7b shows the node degree evolutions for each node separately for sample epilepsy subject. The fluctuation of node degree is shown in the separate subjects from the two groups: epilepsy and control. However, it is hard to draw conclusion about the two groups. That is why we resort to study the group differences using F-tests.

**Fig. 7:**
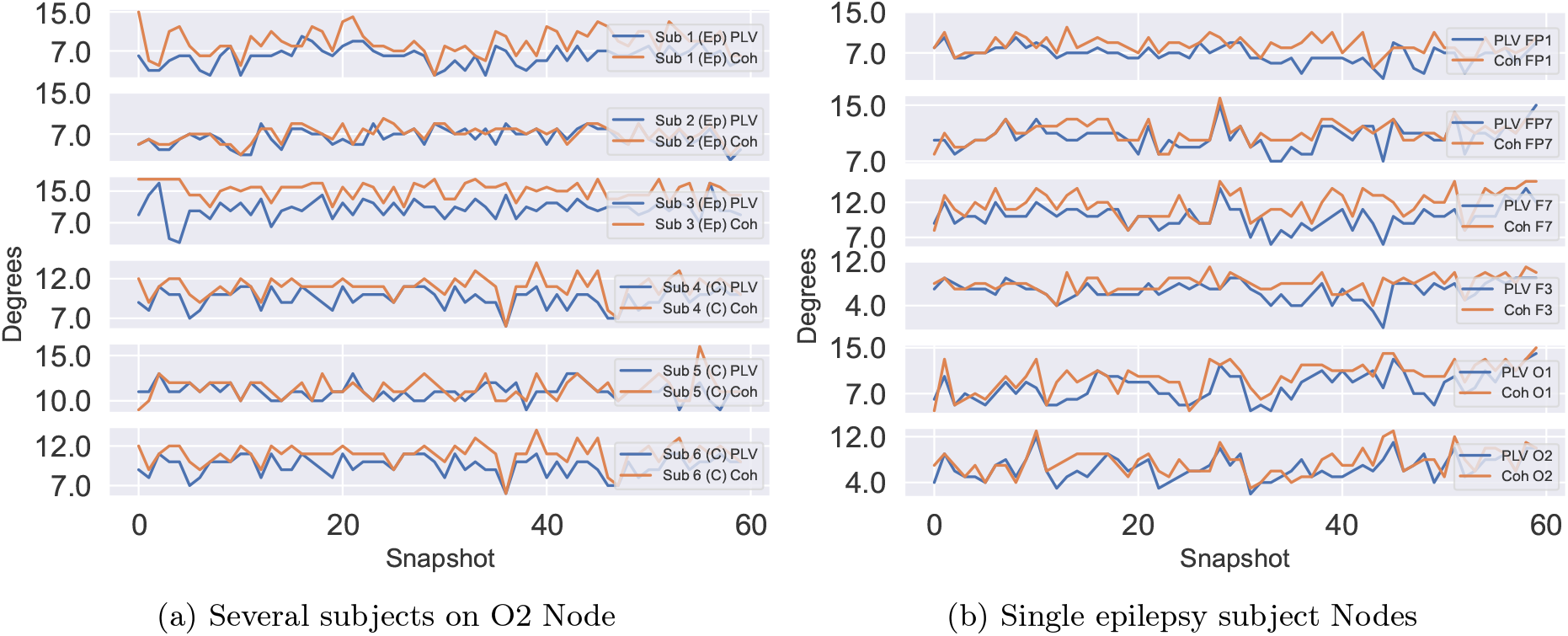
Dynamic changes in node degrees for a) different control (C) and epilepsy (Ep) subjects for one node (O2) using Phase Locking value and Coherence connectivity measures and b) Single epilepsy subject on several nodes

Using the F-test to study differences between the two groups, we noticed that the p-values is 0.045 and 0.0046 for the standard deviations of the node degree evolution for the two groups for the PLV and Coherence, respectively. We reject the null hypothesis. This means that there are significant differences between the two groups. It is shown in figures 8 that healthy control group has the deviation in node degrees are higher than epilepsy group.

**Fig. 8:**
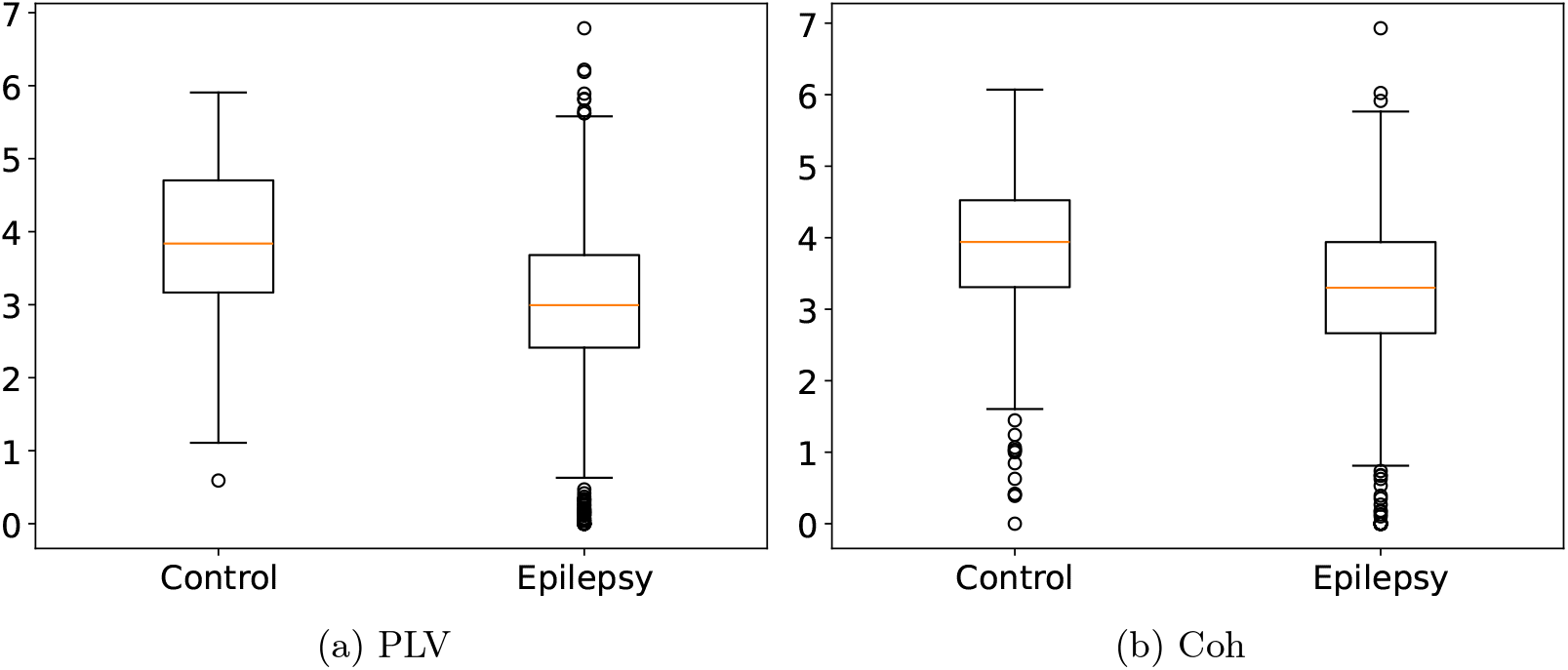
Control vs epilepsy differences in deviation in the node degree evolution over time.

Similarly, the F-test is used to study the differences between the two groups, we noticed that the p-values is 0.000014 and 0.00075 for the mean at the node degree for the two group for the PLV and Coherence, respectively. It is shown in figure 9 that healthy control group has the mean in node degrees that is higher than epilepsy group.

**Fig. 9:**
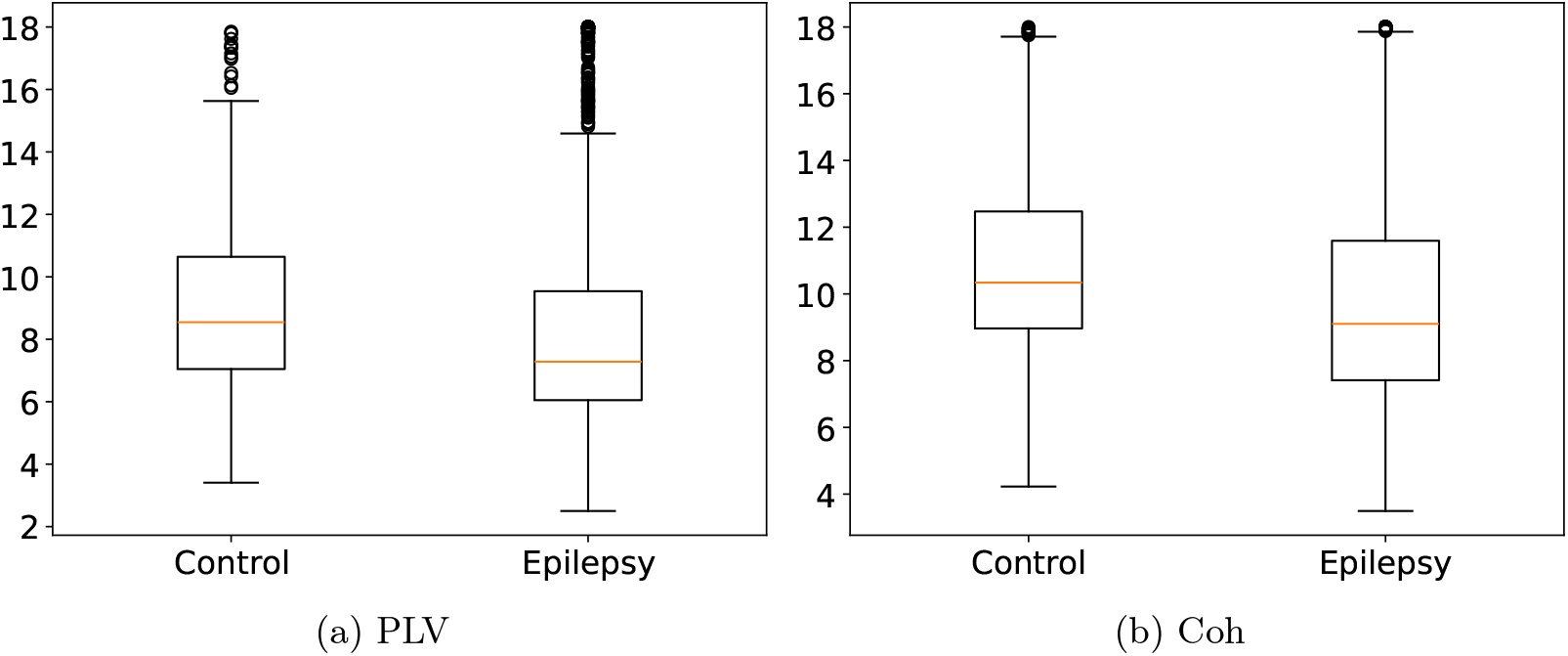
Control vs epilepsy differences in means of the node degrees.

Furthermore, F-Tests are used to signify the difference between the two groups at the node level. Starting with PLV connectivity measure, we notice that there are no significant difference between the two groups using the time fluctuations of node degrees. However, several nodes show significant difference between the two groups when using the mean of node degrees. Those nodes are O2, O1, T5, F8, Pz and Cz as shown in figure 10. On the other hand, for the Coherence connectivity, we notice that there are significant difference in both at the nodes level and the mean of the time fluctuations of the two groups as shown in 10.

**Fig. 10:**
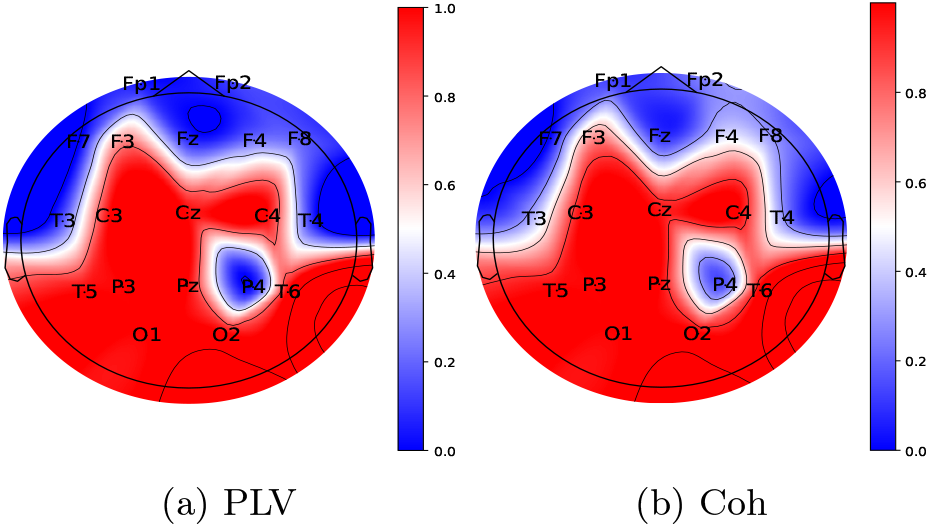
Node degree mean significance obtained by 1-p-value for the a) Phase Locking value (PLV) and b) Coherence (Coh) connectivity measures.

### 3.3 Graph Neural Networks (GNNs)

For further analysis of the dynamic connectivity, we run experiments as explained earlier in section 2.2 using the GNNs over the three EEG datasets and report the results in this section. The used architectures have number of parameters between 35, 199 and 57, 175 as shown in table 1 depending on the used graph operator.

**Table 1:** Number of parameters of the used network for each temporal operator.

| Operator | Number of Parameters |
| --- | --- |
| A3T-GCN | 35199 |
| GCLSTM | 38903 |
| TGCN | 34999 |
| DCRNN | 57175 |
| GConvGRU | 40375 |

Table 2 reports the prediction accuracies and F1 scores for all experimental datasets. The table reports the metrics using the different graph temporal operators on the three used datasets. In the TUAB dataset, All the operators show comprabale performances. The operators AT3GCN and GCLSTM, GConvGRU operators achieved the best prediction accuracy and F1 score using the Coherence and Phase Locking Values as inputs by small margins. In the TUEP dataset, AT3GCN the best scores using the Coherence and Phase Locking Values as inputs. In the NMT dataset, AT3GCN achieved the best scores over all other operators by larger margins.

**Table 2:**
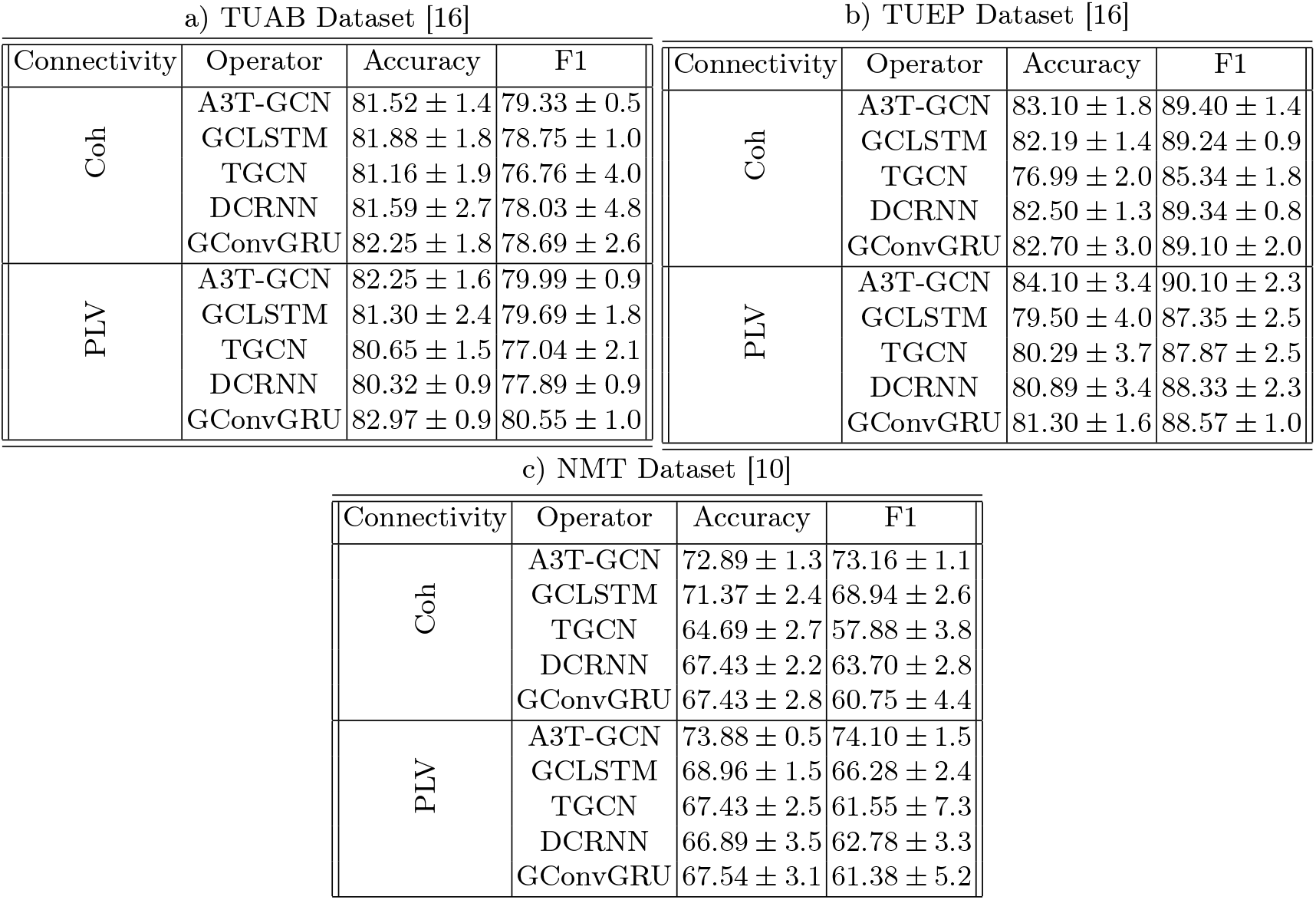
Different temporal graph operators performances on different EEG datasets with Coherence (Coh) and Phase locking Value (PLV) connectivity measures.

| a) TUAB Dataset [16] |  |  |  | b) TUEP Dataset [16] |  |  |  |
| --- | --- | --- | --- | --- | --- | --- | --- |
| Connectivity | Operator | Accuracy | F1 | Connectivity | Operator | Accuracy | F1 |
| Coh | A3T-GCN | $81.52 \pm 1.4$ | $79.33 \pm 0.5$ | Coh | A3T-GCN | $83.10 \pm 1.8$ | $89.40 \pm 1.4$ |
| | GCLSTM | $81.88 \pm 1.8$ | $78.75 \pm 1.0$ | | GCLSTM | $82.19 \pm 1.4$ | $89.24 \pm 0.9$ |
| | TGCN | $81.16 \pm 1.9$ | $76.76 \pm 4.0$ | | TGCN | $76.99 \pm 2.0$ | $85.34 \pm 1.8$ |
| | DCRNN | $81.59 \pm 2.7$ | $78.03 \pm 4.8$ | | DCRNN | $82.50 \pm 1.3$ | $89.34 \pm 0.8$ |
| | GConvGRU | $82.25 \pm 1.8$ | $78.69 \pm 2.6$ | | GConvGRU | $82.70 \pm 3.0$ | $89.10 \pm 2.0$ |
| PLV | A3T-GCN | $82.25 \pm 1.6$ | $79.99 \pm 0.9$ | PLV | A3T-GCN | $84.10 \pm 3.4$ | $90.10 \pm 2.3$ |
| | GCLSTM | $81.30 \pm 2.4$ | $79.69 \pm 1.8$ | | GCLSTM | $79.50 \pm 4.0$ | $87.35 \pm 2.5$ |
| | TGCN | $80.65 \pm 1.5$ | $77.04 \pm 2.1$ | | TGCN | $80.29 \pm 3.7$ | $87.87 \pm 2.5$ |
| | DCRNN | $80.32 \pm 0.9$ | $77.89 \pm 0.9$ | | DCRNN | $80.89 \pm 3.4$ | $88.33 \pm 2.3$ |
| | GConvGRU | $82.97 \pm 0.9$ | $80.55 \pm 1.0$ | | GConvGRU | $81.30 \pm 1.6$ | $88.57 \pm 1.0$ |
| c) NMT Dataset [10] |  |  |  |  |  |  |  |
| Connectivity | Operator | Accuracy | F1 |  |  |  |  |
| Coh | A3T-GCN | $72.89 \pm 1.3$ | $73.16 \pm 1.1$ | | | | |
| | GCLSTM | $71.37 \pm 2.4$ | $68.94 \pm 2.6$ | | | | |
| | TGCN | $64.69 \pm 2.7$ | $57.88 \pm 3.8$ | | | | |
| | DCRNN | $67.43 \pm 2.2$ | $63.70 \pm 2.8$ | | | | |
| | GConvGRU | $67.43 \pm 2.8$ | $60.75 \pm 4.4$ | | | | |
| PLV | A3T-GCN | $73.88 \pm 0.5$ | $74.10 \pm 1.5$ | | | | |
| | GCLSTM | $68.96 \pm 1.5$ | $66.28 \pm 2.4$ | | | | |
| | TGCN | $67.43 \pm 2.5$ | $61.55 \pm 7.3$ | | | | |
| | DCRNN | $66.89 \pm 3.5$ | $62.78 \pm 3.3$ | | | | |
| | GConvGRU | $67.54 \pm 3.1$ | $61.38 \pm 5.2$ | | | | |

For further validation, table 3 shows the comparison with several baseline graph models on the TUAB dataset for fair comparison. The proposed models in this study outperforms other graph models in the same data except the F1 score by the model GATs+LSTM variant that was proposed by [29].

**Table 3:** Comparison between different baseline graph models on TUAB dataset [16].

| Model | Accuracy | F1 |
| --- | --- | --- |
| InstaGATs [5] | $77.25 \pm 1.8$ | $76.83 \pm 1.6$ |
| GNN [5] | $74.01 \pm 1.2$ | $73.40 \pm 1.7$ |
| GATs+LSTM 1 [29] | 76.92 | 77.08 |
| GATs+LSTM 2 [29] | 81.27 | 82.93 |
| STAGIN-EEG [27] | 78.91 | 77.31 |
| GConvGRU with Coh (this study) | $82.25 \pm 1.8$ | $78.69 \pm 2.6$ |
| GConvGRU with PLV (this study) | $82.97 \pm 0.9$ | $80.55 \pm 1.0$ |
| AT3GCN with Coh (this study) | $81.52 \pm 1.4$ | $79.33 \pm 0.5$ |
| AT3GCN with PLV (this study) | $82.25 \pm 1.6$ | $79.99 \pm 0.9$ |

## 4 Discussion

In brain connectivity analysis, the degree of a node or brain region indicates how many other regions it is functionally or structurally connected to. Abnormal networks and node degree may suggest disruptions in network communication, which can be associated with neurological disorders. In this study, we used network efficiency, node degree and more advanced GNNs methods for this task.

### 4.1 Network Efficiency

From the study of network efficiency, It is noted that the local and global efficiency of two groups: healthy and Epileptic patients are different. It is noted that efficiency is lower in epilepsy patients compared to healthy control. This is noted in both global and local efficiencies. This indicates reduced connection strength in the network. This is aligned with previous research such as [25], [13]. Global efficiency captures the ability of the network to integrate information globally. High global efficiency implies that a certain node can be reached from any other node through relatively short paths, indicating an efficient global communication structure. Local efficiency reflects how well the network preserves communication within a local neighborhood if a particular node fails. Higher local efficiency implies a more robust and clustered local structure.

The global and local network efficiency fluctuations are not straightforward to understand. The F-tests indicates that local efficiency fluctuations are significant in the two groups. On the other hand, global efficiency fluctuations are not significant. This fluctuations in local efficiency in the two subjects groups means that the local neighbourhood communications can be different between the two subject groups. The local network efficiency fluctuations abnormality can be correlated with epilepsy. This argument will confirmed in the node degree analysis.

### 4.2 Node Degree

In connection with network efficiency, Node degree is related as the edges with lower strength connections are excluded in the networks. This leads generally to lower node in epilepsy subjects in comparison with healthy group in the TUEP dataset. Node degree evolution differences has shown to be significant in the two used groups. The node degree evolution deviation from the mean is higher in healthy than Epileptic group.

Both fluctuation and mean have shown to be significant in between the two groups of healthy than Epileptic group. Additionally, there are significant nodes to show the fluctuations between deviations of the two groups. Several nodes show significant difference between the two groups when using the mean of node degrees. Those nodes are from the Frontal, Occipital, Parietal and temporal cortex.

There are several research in literature that highlight that nodes within or near the Seizure onset zone tend to have higher node degrees, reflecting increased activity and connectivity in those areas. In the same time, some regions may exhibit lower node degrees, indicating reduced communication and possibly disrupted network function. [24], [25], [3], [8].

### 4.3 Graph Neural Networks

The proposed models using the different graph operators outperform or provides comparable performances to the several baseline graph models. This indicates the usefulness of the temporal dynamics of connectivity in the analysis of EEG. The Attention Temporal Graph Convolutional Network (AT3-GCN) consistently achieved the best performances in the three used datasets. This suggests the effectiveness of attention to capture global variation in EEG.

Using the TUAB dataset, there are several trials using other model architectures based on Deep CNN such as [18] and [20] which achieved superior performances. But our objective was to compare graph neural networks baselines for fair comparison. Additionally, working with EEG data involves extensive preprocessing and highly controlled experiment setting which may make the comparison less fair.

## 5 Conclusion and Future Directions

In this study, we explored the significance of dynamically evolving functional connectivity in the training of Graph Neural Networks (GNNs). We also employed F-tests to statistically assess changes in network-level features. Our proposed GNN models offer a lightweight and parameter-efficient baseline while achieving competitive performance against previous GNN approaches.

By modeling dynamic brain networks, our approach captures temporal variations that are often overlooked by static GNN baselines. This dynamic perspective offers a more realistic representation of brain function. In the future, incorporating self-supervised learning approaches could help extract more meaningful embeddings, further enhancing classification performance. Moreover, the model could be extended to perform forecasting of future brain network states, which may aid in understanding brain dynamics and pathological transitions, particularly when paired with explainable machine learning techniques.

An ablation study on model architectures and feature sets is also recommended to assess their relative contributions. This analysis would provide deeper insight into the model’s predictive behavior, revealing which features or architectural components drive classification decisions. Such analysis can be further supported using interpretable ML methods to identify discriminative brain regions and connectivity patterns.

Our approach to dynamic connectivity was also evaluated on the TUEP epilepsy dataset. While our primary goal was to demonstrate the method’s utility in pathology-specific classification rather than general abnormality detection, further work is required to fully capture the complexity of epilepsy disorder in deeper details.

Finally, we examined brain network efficiency that reflects the integrative capacity of functional networks and its alteration in pathologies such as epilepsy. These metrics, when combined with node-level analyses like node degree, can yield biologically meaningful insights. Integrating these findings with explainable AI tools may further enhance our understanding of how different brain states contribute to neurological disorders.

## Acknowledgments

This work is part of the AI-Mind project. AI-Mind has received funding from the European Union’s Horizon 2020 research and innovation program (https://www.ai-mind.eu).

